# The intestinal microbiota predisposes to traveller’s diarrhoea and to the carriage of multidrug-resistant Enterobacteriaceae after travelling to tropical regions

**DOI:** 10.1101/265033

**Authors:** Stefano Leo, Vladimir Lazarevic, Nadia Gaïa, Candice Estellat, Myriam Girard, Sophie Matheron, Laurence Armand-Lefèvre, Antoine Andremont, The VOYAG-R Study Group, Antoine Andremont, Laurence Armand-Lefèvre, Olivier Bouchaud, Yacine Boussadia, Pauline Campa, Bruno Coignard, Paul-Henri Consigny, Assiya El Mniai, Marina Esposito-Farèse, Candice Estellat, Pierre-Marie Girard, Catherine Goujon, Isabelle Hoffmann, Guillaume Le Loup, Jean-Christophe Lucet, Sophie Matheron, Nabila Moussa, Marion Perrier, Gilles Pialoux, Pascal Ralaimazava, Etienne Ruppè, Daniel Vittecoq, Ingrid Wieder, Benjamin Wyplosz, Jacques Schrenzel, Etienne Ruppé

## Abstract

The risk of acquisition of multidrug-resistant Enterobacteriaceae (MRE) and of occurrence of diarrhoea is high when travelling to tropical regions. The relationships between these phenomena and the composition of human gut microbiota have not yet been assessed. Here, we investigated the dynamics of changes of metabolically active microbiota by sequencing total RNA from faecal samples taken before and after travel to tropical regions. We found that the occurrence of diarrhoea during the travel was associated with a higher relative abundance of Prevotella copri before departure and after return. The composition of microbiota, before travel as well as at return, was not correlated with the acquisition of MRE. However, the clearance of MRE one month after return was linked to a specific pattern of bacterial species that was also found before and after return.

## Introduction

Multidrug-resistant Enterobacteriaceae (MRE) that produce extended-spectrum beta-lactamases (ESBLs), plasmid-encoded AmpC-type cephalosporinases (pAmpC) and/or carbapenemases (CP) have been spreading in the community over the last two decades [1]. MRE represent a major public health issue, as a limited number of antibiotics remains active against these bacteria while very innovative antibiotics are expected to reach the market in the near future [2]. The spread of MRE has been particularly intense in tropical regions, likely owing to poor hygiene conditions and uncontrolled antibiotic usage [3]. Consequently, between 14% and 69% of travellers to tropical regions are reported to acquire MRE, depending on the cohort and specific destination [4, 5]. Besides, a high proportion of travellers to these destinations also report the occurrence of diarrhoea (traveller’s diarrhoea) during their trip [4-8]. Among travellers who acquire a MRE during a travel to tropical regions, the observed median carriage is short (≤1 month) [5, 6]. Still, some of them experience long-term carriage of MRE, extending up to 1 year in 2.2-11% cases [5, 6].

The capacity of the intestinal microbiota to resist long-term settlement of exogenous bacteria (including MRE) is referred to as colonization resistance [9-11] which is mainly exerted by anaerobes [9, 10]. Antibiotics with high activity against anaerobes strongly affect the capacity of the microbiota to prevent colonization by exogenous microorganisms, and thus favour the acquisition and expansion of resistant bacteria [12]. Hence, the restoration of the intestinal microbiota through faecal material transplantation (FMT) has been shown to lower the intestinal concentrations of vancomycin-resistant enterococci (VRE) [13], MRE [14] and more globally, of antibiotic resistance genes [15]. Moreover, the administration of a limited set of intestinal bacteria to mice colonised with VRE reduced the intestinal densities of VRE, suggesting that specific bacteria are involved in colonization resistance [16].

To date, the link between the composition of the intestinal microbiota and the acquisition of MRE during travel and their clearance after return has not been assessed.

Here, we questioned whether the composition of the intestinal microbiota could be associated with the occurrence of diarrhoea and the acquisition of MRE during travel and to the clearance of MRE after return.

## Methods

### Population

The travellers cohort originates from the VOYAG-R study (ClinicalTrials.gov n°NCT01526187) funded by the French Ministry of Health [6, 17]. From February 2012 to April 2013, subjects attending six international vaccination centres in the Paris area, prior to traveling to tropical regions, were asked to provide faecal samples before and after their trip. Only volunteers who had no detectable MRE in the faeces taken before their departure, were asked to send a further stool sample within a week after their return. Travellers who revealed positive for MRE after their return were asked to provide additional stool samples 1, 2, 3, 6 and 12 months after their return until MRE was no longer detected. Among the 574 included subjects, 292 (51%) acquired at least one MRE [6]. For the present analysis, a specific sub-cohort of 43 subjects (15 from Sub-Saharan Africa, 13 from Latin America and 15 from Asia) were chosen. No particular criteria of inclusion were applied except the fact that we wanted to have almost the same number of travellers per tropical region. In addition to a native stool sample, volunteers were asked to ship a stool aliquot diluted in another 200 mL vial containing 50 mL of RNAlater^®^ solution (Applied Biosystems, Villebon-sur-Yvette, France) for metagenomic analysis. In this sub-cohort of 43 travellers, 18 subjects (41.8%) acquired an MRE (mostly ESBL-producing *Escherichia coli*) during their trip. All of them provided a stool sample one month after they returned, among whom 6 were still carrying a MRE. Thus, 104 samples were selected (43 before travel, 43 at return and 18 one month after travel).

### RNA extraction from stool samples

Total RNA (mostly made of ribosomal RNA *e.g.* 16S, 23S and 5S) was extracted for 104 stool samples using the RNeasy Plus Mini Kit (Qiagen, Gaithersburg, USA). The concentration of RNA obtained was measured by Qubit RNA BR Assay Kit or Qubit RNA HS Assay Kit (Life Technologies, Reinach, Switzerland). For simplicity, the terms 16S and 23S refer to 16S and 23S ribosomal RNAs, respectively. The integrity of RNA (ratio 16S/23S) was assessed by the RNA 6000 Nano Kit and RNA 6000 Pico Kit (Agilent Technologies, Plan-les-Ouates, Switzerland) on the Bioanalyzer 2100 system (Agilent Technologies, Waldbronn, Germany). The detailed protocol can be found in the Supplementary Methods.

### Sequencing and reads processing

One hundred ng of purified RNA in 10 µL total volume were sent to LGC (Berlin, Germany) for (*i*) cDNA synthesis using NEBNext mRNA First/Second Strand Synthesis Module (New England Biolabs, Ipswich, USA) (*ii*) shotgun library preparation using NuGEN Ovation Rapid Library System (NuGEN, San Carlos, USA), and (*iii*) sequencing (2 × 300bp) of size-selected fragments (about 300-400 bp) using v3 Illumina chemistry and half the capacity of a MiSeq (Illumina, San Diego, USA) flow cell. The paired reads were merged using BBMerge 35.43 (http://bbmap.sourceforge.net), trimmed and quality-filtered by Mothur v1.35.1 [18]. The average number of merged quality-filtered pairs exploitable for mapping was 89,351 (median = 87,728) per sample, the average length of reads after merging was 355 (range = 200–560). Quality-filtered merged reads were mapped onto the 16S Greengenes operational taxonomic units (OTUs) database pre-clustered at 97% identity [19] with USEARCH [20] by considering an identity of ≥97% and ≥97% query coverage (see Supplementary Methods). Downstream analyses were carried out using the taxonomic information of the matching full-length reference 16S gene sequences. In parallel, we aligned quality-filtered merged reads against the SILVA ribosomal 23S/28S database [21] with USEARCH using the same settings as for the 16S profiling. Then, we selected the reads that mapped to bacterial 23S genes but had no hit against eukaryotic 28S sequences. Reads classified to the same taxon were summed up. The mapping rates to 16S and 23S genes are reported in Supplementary Methods. The relative abundance of a given OTU or taxon was computed as percentage of the total number of reads in a sample, mapping to that OTU or taxon.

### Bio-statistical analyses

The variables used in this study are detailed in the Supplementary Methods. To assess similarities and differences between bacterial communities in response to predictor variables, we performed principal coordinates analysis (PCoA) either in PRIMER v6 (PRIMER-E Ltd, Plymouth, UK) or in the R software v3.2.3 with the vegan v2.3-5 and ade4 v1.7-4 packages. Permutational analysis of variance (global and pairwise PERMANOVA, with 9,999 permutations, unrestricted permutation of raw data and Type III sums of squares) and distance-based linear models (DistLM, with 9999 permutations) based on the Bray-Curtis similarity matrices of square-root transformed OTUs’ or taxa relative abundance were carried out in PRIMER. Ecological indices were determined after rarefying the 16S data set to the lowest number of reads assigned to the OTUs in any of the samples (10,465). Observed richness (S) and Shannon diversity were calculated in PRIMER.

Taxa with significantly different relative abundances between two conditions were identified in R software using the Wilcoxon rank sum test or Wilcoxon signed rank test (with a significance threshold of p<0.05 without adjustments for multiple comparisons [22]). Besides, OTUs contributing to discrimination between groups of samples were also identified using similarity percentage analysis (SIMPER, PRIMER-E) and the on-line tool Linear discriminant analysis Effect Size (LeFSe) [23].

### De novo 16S gene assembly

Samples taken before departure from 39 subjects who were not treated with antibiotics during the travel were considered for this analysis. First, reads mapping to a given OTU were collected from each sample and assembled with CAP3 (version date: 02/10/15) [24] using default settings. Then, the 16S genes assembled from each sample and corresponding to the same OTU were aligned and the consensus sequence was determined again with CAP3. Finally, the taxonomic assignment of the obtained consensus sequence of 16S genes was done by BLASTing [25] against the non-redundant 16S SILVA and Greengenes databases using a minimum percentage identity of 97. Hits with the highest percentage of identity and lowest e-value of alignment were retained. In case of multiple choices with equal scores, the hit with the lowest taxonomic rank was selected.

## Results

### Effect of travelling on microbiota composition

To test the hypothesis whether the modalities and the duration of travelling to tropical regions has an impact on microbiota composition, we analysed the microbiota profiles at return of 39 travellers that were not administered antibiotics (except doxycycline for malaria prophylaxis) during their trip. We did not observe any significant influence on microbiota composition in relation to the visited region, the type of travel, duration of travel, to the use and the type of malaria prophylaxis (Table 1). However, the proportions of Enterobacteriaceae increased during the travel in all the 39 individuals (Supplementary Figure 5).

**Table 1.** Summary of global PERMANOVA analyses performed on the cohort of 39 travellers at return.

| Variables | Conditions | rRNA subunit | Pseudo-F | p-value |
| --- | --- | --- | --- | --- |
| Region | Sub-Saharan Africa / Latin America / Asia | 16S | 0.89 | 0.75 |
|  |  | 23S | 0.81 | 0.82 |
| Type of travel | Organised tour/ Family / Mix of all-inclusive resorts and organised tours / Backpacking / All-inclusive resort | 16S | 1.08 | 0.25 |
|  |  | 23S | 1.11 | 0.23 |
| Duration of travel (*) | From 1 to 9 weeks | 16S | 1.04 | 0.35 |
|  |  | 23S | 0.82 | 0.71 |
| Malaria prophylaxis | No / Yes | 16S | 0.79 | 0.86 |
|  |  | 23S | 0.76 | 0.80 |
| Type of malaria prophylaxis | Atovaquone-proguanil / Chloroquine / Doxycycline / None | 16S | 0.95 | 0.57 |
|  |  | 23S | 0.92 | 0.63 |

(\*) The test done was DistML (see Methods).

Moreover, we analysed microbiota profiles before departure, at return and one month after the return of 17 travellers who acquired MRE during their journey and were not treated with antibiotics. We observed that the faecal samples taken at different time points (before departure, at return and one month after return) clustered by subject (global PERMANOVA test, p-value <0.0001; Supplementary Figure 12), while travel did not have a significant impact on microbiota profiles (pairwise PERMANOVA tests, p-values ranged between 0.3-0.9).

### Occurrence of diarrhoea during the travel

In pre-travel microbiota of people who suffered from diarrhoea during the travel (n=9) the relative abundance of *Prevotella copri* species was >2-fold higher than in subjects who did not experience this condition (n=34) (Wilcoxon rank sum test, p-value <0.05; Supplementary Figure 1C-D). Nevertheless, the overall composition of pre-travel microbiota was not significantly associated with the occurrence of diarrhoea during travel (Table 2; Supplementary Figure 1A).

**Table 2.** Summary of pairwise PERMANOVA tests performed at each travel time point*.

| Travel time point | Number of travellers | Variable | rRNA | t | p-value |
| --- | --- | --- | --- | --- | --- |
| Before travel | 43 | Diarrhoea during travel | 16S | 1.05 | 0.25 |
|  |  |  | 23S | 1.1 | 0.18 |
|  |  | MRE acquisition | 16S | 1.09 | 0.15 |
|  |  |  | 23S | 1.05 | 0.29 |
| At return | 39 | Diarrhoea during travel | 16S | 1.44 | 0.003 |
|  |  |  | 23S | 1.8 | 0.0007 |
|  |  | MRE acquisition | 16S | 1.12 | 0.12 |
|  |  |  | 23S | 1.13 | 0.15 |
| One month after the return | 17 | Diarrhoea during travel | 16S | 1.04 | 0.3 |
|  |  |  | 23S | 0.97 | 0.6 |
|  |  | MRE carriage at return | 16S | 1.21 | 0.01 |
|  |  |  | 23S | 1.16 | 0.07 |

The number of samples analysed varied among travel time points according to the use of antibiotics and MRE acquisition. Before travel we have considered all the 43 travellers. At return we analysed only subjects who did not take antibiotics during the travel. One month after the return, we considered travellers who acquired MRE during travel and were not treated with antibiotics after the trip.

We, then, analysed the intestinal microbiota profiles at return by excluding those 4 travellers who had taken antibiotics during their trip. Microbiota profiles from subjects who had diarrhoea significantly differed from those who had not (Tables 2; Figure 1A). The occurrence of diarrhoea was associated with a microbiota significantly less rich and diverse at return as compared to the microbiota of individuals who did not experience this condition during the travel (Wilcoxon rank sum test, p-value <0.05; Figure 1B).

**Figure 1.**
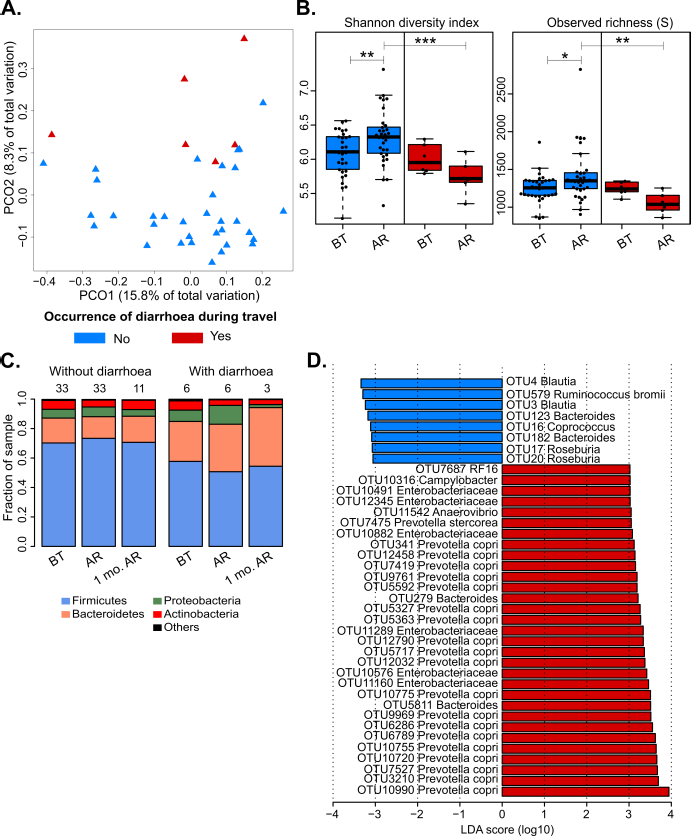
Microbiota changes with respect to the occurrence of diarrhoea during the travel. (A) PCoA plot showing the distribution of samples taken at the return, from travellers who had (red triangles) or not (blue triangles) diarrhoea during the travel. Percentage of variation explained by the first two axes is indicated. (B) Boxplots and dot plots depicting the values and their corresponding distributions of ecological indexes, computed at the 16S OTU level, before travel (BT) and at return (AR), between travellers experiencing or not diarrhoea. Stars correspond to the following p-values: * = <0.05; ** = <0.01; *** = <0.001. (C) Bar plots reporting the averaged relative abundance (/100) of phyla detected by mapping to 16S Greengenes database in travellers without diarrhoea (n=33 of which 11 were still MRE positive one month after the return) and those with (n=6 of which 3 were still MRE positive one month after the return). Numbers at the top of the bars indicate the amount of samples for each travel time point/condition. (D) Bar plot reporting log10-transformed LDA scores of OTUs selected by LEfSe analyses (p-value <0.05). Cohorts of samples and colour label are the same as explained in (A). Only OTUs with a log10 LDA score of at least 3 are represented. Greengenes taxonomy identifiers for all OTUs are reported in Supplementary Table 3.

At return, people having experienced diarrhoea during their travel presented increased proportions of Bacteroidetes and Proteobacteria phyla and decreased proportions of Firmicutes phylum compared to travellers without diarrhoea (Figure 1C). The great majority of OTUs found to be differentially abundant after travel between people with and without diarrhoea mapped to Enterobacteriaceae family, Clostridiales order and the *P. copri* species (LEfSe and Wilcoxon rank sum test, p-value < 0.05; Figure 1D; Supplementary Figures 2A, 5, 6, 7). In particular, the faecal samples taken at return from travellers who reported diarrhoea showed a >2-fold higher proportion of Enterobacteriaceae and *Prevotella copri* and a <2-fold lower proportion of Clostridiales as compared to travellers without diarrhoea. The relative abundance of *P. copri* was increased at return only in a fraction of travellers reporting diarrhoea (Supplementary Figure 6).

The results obtained for the samples collected at return from travel were also confirmed by 23S analyses (Supplementary Figure 2B-C-D-E). Most Enterobacteriaceae members associated with diarrhoea mapped to 23S genes of *Escherichia-Shigella,* while in the 16S analysis the discriminating OTUs from Enterobacteriaceae were not classified at the genus level. Clostridiales included genera belonging to Ruminococcaceae *(Ruminococcus),* Bacteroidaceae (*Bacteroides*) and Lachnospiraceae *(Roseburia;* only for 16S dataset: *Lachnospira and Blautia).*

We then investigated whether diarrhoea had protracted effects on gut bacteria composition one month after the return. Therefore, we analysed samples from 17 travellers who did not take antibiotics during the first month following the return. We did not detect significant differences in the overall microbiota composition in relation to diarrhoea (Table 2, Supplementary Figure 8A). Enterobacteriaceae relative abundance returned close to basal level in all 17 travellers but with statistical significance only when the analyses included the individuals who did not suffer from diarrhoea (Wilcoxon signed rank test, p-value <0.05, Supplementary Figure 9). One month after return, *Prevotella copri* abundance was still increased in travellers who had diarrhoea (Supplementary Figure 6) whereas Clostridiales members were more abundant in individuals without diarrhoea (Supplementary Figure 7).

### Acquisition and carriage of MRE

The acquisition of MRE was not associated with a specific microbiota profile neither before travel nor at return (Table 2; Supplementary Figures 1B, 10).

We also investigated the association of microbiota composition and the MRE carriage one month after return in the 17 travellers found MRE-positive at return and not treated with antibiotics. In this case, the composition of the intestinal microbiota of the subjects whose samples became MRE-negative (n=11) was significantly different from those whose samples remained MRE-positive (n=6) (Table 2). OTUs assigned to *Bifidobacterium adolescentis* (OTU6129) and to some Clostridiales (OTU61, OTU7384, OTU405, OTU8089) as well as *Bifidobacterium* strains detected by 23S analyses, were enriched in individuals who had cleared MRE one month after the return (LEfSe and Wilcoxon rank sum test, p-value <0.05; Figure 2A-B; Supplementary Figure 11A-B). Already before departure, the proportions of these bacteria were higher in MRE carriers who cleared their carriage one month after return than in those who were still positive Figure 2A; Supplementary Figure 11A). For most OTUs associated with MRE clearance, the species level taxonomy of *de novo* assembled 16S gene was not available (Supplementary Table 1).

**Figure 2.**
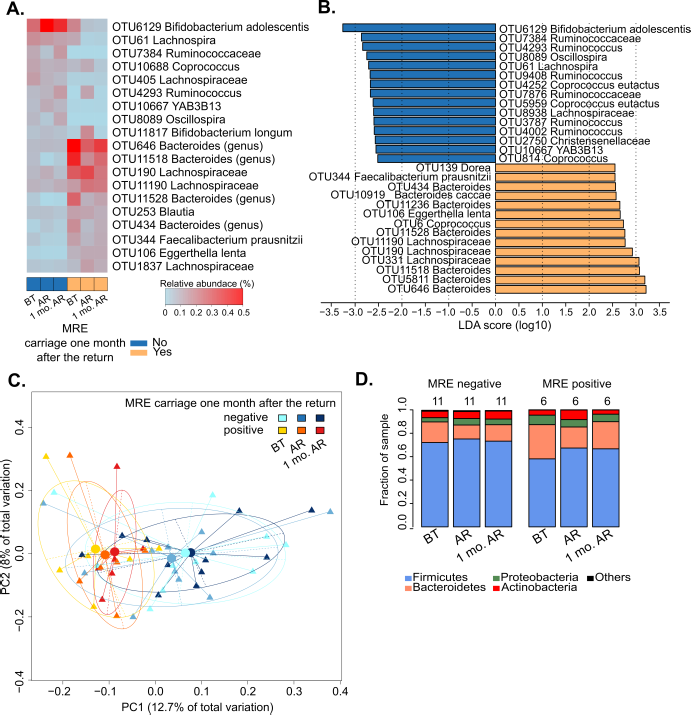
Comparison of travellers who acquired MRE and remained positive (n=11) or became negative (n=6) one month after return. (A) Heat map illustrating the mean relative abundance (expressed as percentage of total reads) of 19 OTUs in the three sampling points (BT, AR and 1 mo. AR = before travel, at return and one month after the return, respectively) of travellers MRE positive (yellow bottom bar) and MRE negative (blue bottom bar) one month after the return. OTUs were selected if the p-value was < 0.05 (Wilcoxon rank sum test), the fold change was of at least 2, and when the mean relative abundance was of at least 0.1% in one of the six represented conditions represented (i.e. BT, AR and 1 mo. AR of MRE-positives and of MRE-negatives one month after the return). Greengenes taxonomy identifiers for all OTUs are reported in Supplementary Table 3. (B) Barplot reporting log10-transformed LDA scores from each OTUs resulting significant (p-value < 0.05) from LEfSe analyses on samples from travellers MRE positive (yellow) and MRE negative (blue) one month after the return. Only OTUs with a log10 LDA score of at least 2.5 were retained for graphic representation. (C) PCoA plot showing the distribution of samples from time points of travellers MRE negative and MRE positive one month after return. PCO1 (12.7%) and PCO2 (8%) represent the degrees of variance of each axis. Samples are grouped in clusters delimited by ellipses. Represented centroids (spheres) capture the origin of each ellipsis. (D) Bar plots reporting the averaged relative abundance (/100) of phyla detected by mapping to 16S Greengenes database in travellers MRE negative (n=11) and MRE positive travellers (n=6) one month after return.

Remarkably, within the subcohorts of 17 travellers, we found that the microbiota profiles of MRE cleared individuals were significantly different from persistent MRE carriers before departure and one month after return (Supplementary Table 2; Figure 2C; Supplementary Figure 11C).

The observed species richness measured before travel was significantly lower in individuals who acquired MRE during the travel and remained MRE carriers one month after return than in those who cleared their carriage (Supplementary Figure 8B). We found that the relative abundance of several OTUs and strains/species mapping to Enterobacteriaceae significantly decreased more than 2 fold one month after the return, independently of MRE carriage (Supplementary Figures 13). Despite that, the abundance of Proteobacteria phylum, to which Enterobacteriaceae belong, did not change over time or in relation to MRE carriage measured one month after the return (Figure 2D; Supplementary Figure 11D).

## Discussion

One of the main results of this study is that travellers who experienced diarrhoea (regardless of the aetiology) had higher intestinal abundance of *P. copri,* before the travel, at return, and one month after, than those who did not. This suggests that subjects with higher relative abundance of *P. copri* could be at higher risk for diarrhoea during travel. Interestingly, *P. copri* has popped up in various metagenomic studies as either a beneficial or pathogenic bacterium: it has been associated to rheumatoid arthritis [26, 27], insulin resistance [28] and parasitic diarrhoea in children [29], but also to good health status [30, 31] and to an improved glucose homeostasis [32]. These contradictory findings could be explained by the high inter-strain and inter-individual genetic variations of this species [33], suggesting that different strains have various functions, antigens and/or metabolites being either beneficial or deleterious for the host. Another characteristic of microbiota profiles of people suffering diarrhoea is decreased species richness at return. Diminution of species richness upon diarrhoea occurrence is also described in one study conducting by 16S profiling the intestinal microbiota of travellers returning to USA from Central America or India [34]. Oppositely, subjects suffering from a Norovirus-caused diarrhoea had a surprisingly higher intestinal richness and diversity than diarrhoea-free travellers. However, in this study low numbers of reads per sample (<3000) were analysed and pre-travel samples were not assessed [34].

We observed that the travellers who did not experience diarrhoea during travel had a microbiota profile enriched in members of *Ruminococcus, Coprococcus,* and *Dorea.* Consistently with our findings, these genera were depleted in 16S microbiota profiles of individuals suffering from post-transplantation [35] and nosocomial diarrhoea, including *Clostridium difficile* infection [36]. Moreover, *Roseburia* and *Ruminococcus* species have been shown to prevent gut inflammation by strengthening gut barrier function in mice [37] and by enhancing starch fermentation in humans [38], respectively. Subjects who did not experience diarrhoea during the travel had a richer and more diverse microbiota after travel than before. However, one month after return, the diversity and richness for those individuals tended to decrease to a baseline (pre-travel) level. Increase in richness and diversity at return when diarrhoea is not experience could reflect the ingestion of new bacteria that are not met in France, and/or the consumption of food that could act as prebiotics for bacteria in the pre-travel sample.

Besides, we observed an increase of Enterobacteriaceae in all travellers, which was more significant in those subjects who experienced diarrhoea during their trip. This could be expected as several diarrhoea agents - *E. coli, Shigella* and *Salmonella* - belong to the Enterobacteriaceae family [39]. This increase was transient and the relative abundance of Enterobacteriaceae returned close to baseline one month after travel.

Another significant result of this study is the association between intestinal microbiota composition and clearance of MRE in healthy travellers. Indeed, we observed that a set of bacteria from the Clostridiales order was more abundant in travellers who had no detectable MRE one month after return than in those who remained MRE carriers. Moreover, this pattern was already observed before travel. In addition, among the subjects who acquired MRE during travel, bacterial richness and diversity at baseline (before travel) were lower in individuals who remained MRE carriers one month after travel than in those who did not. Altogether, our findings support the concept that the intestinal microbiota before travel may not predispose to the acquisition of MRE, but will affect its clearance in case of acquisition. These results are in line with the observations performed in mice in which some specific OTUs were shown to be associated with the clearance of vancomycin-resistant enterococci [16] - *Listeria monocytogenes* [40] and *Clostridium difficile* [41] - but also with the MRE clearance after faecal material transplantation [14]. Nonetheless, these phenomena were observed after an alteration of the microbiota by antibiotics, whereas our observations were obtained from healthy, antibiotic-free travellers.

The main limitation of this study is that the number of samples is relatively low with regards to some variables such as MRE carriage after the return and the occurrence of diarrhoea. Nonetheless, we combined several bio-statistical and bioinformatics approaches which produced consistent results and therefore strengthen our findings. Of note, we took advantage of using total RNA (rich in ribosomal RNA) to bypass the need for amplification of a specific region of the 16S gene (that leads to biases) and to allow a separate analysis on 16S and 23S taxonomic markers. Hence, despite the low number of samples in some sub-groups, we are confident with our conclusions because of the consistency of the results throughout the various bio-statistical and bioinformatic analyses performed. Another limitation is that we could not precisely identify the bacterial species associated with the intestinal clearance of MRE despite our attempts to assemble the full 16S genes, hindering the realization of in vitro and in vivo experiments to demonstrate the precise role of these bacteria. Metagenomic sequencing that allows functional analysis could now be used on a similar setting to identify the genes associated to the clearance of MRE irrespectively of the taxonomy of their host. Finally, the aetiological agents of traveller’s diarrhoea were not sought as it was outside of the VOYAG-R study’s scope. Consequently, we could not link the composition of the microbiota to the presence of a specific pathogen. Still, the intestinal alteration due to traveller’s diarrhoea seems to be pathogen-independent [34].

In conclusion, we showed that the composition of intestinal microbiota is associated with the risk of occurrence of diarrhoea in healthy travellers and of carriage of MRE in those who acquired MRE during the travel. Our results call for further functional explorations of the interplay between the intestinal microbiota, traveller’s diarrhoea and MRE carriage.

## Data Availability

Mothur-quality-filtered sequences were deposited as fastq files at the European Nucleotide Archive (ENA) under the project PRJEB24843. Prior to sequences’ submission, reads assigned to human genome (vGRCh38.p10) by CLARK (v1.2.3.2) [42] were removed. CLARK taxonomic classification was performed at the phylum level (Chordata).

## Acknowledgements

We are grateful to Dr María-José Gosalbes (Área Genómica y Salud, FISABIO-Salud pública, Valencia, Spain), Pr Andrés Moya (Cavanilles Institute on Biodiversity and Evolutionary Biology, University of Valencia, Spain) and to Anaïs Gondoin (SATT Ile de France, Paris, France) for their assistance in this project. Again, we thank all the travellers, who agreed to participate in this study, and the personnel of the international vaccination centers.

## Author contributions

ER, LAL and AA conceived and designed the study. ER, JS and VL supervised the study. The VOYAG-R study group performed the princeps’s study. MG and SL performed RNA extraction. E.R., V.L., S.L. designed the statistical analyses. SL and ER analysed the data. NG helped with read pre-processing. SL, VL and ER wrote the manuscript. SL assembled all the figures. CE and SM revised the manuscript. All authors read and approved the final manuscript.

